# The splicing kinase PRPF-4 is required for somatic development and germline function in *C. elegans*

**DOI:** 10.64898/2026.08.28.747746

**Authors:** Willie C. Barron, Xiaolu Wei, Sakia Ferdousy, Lishuang Zhu, Fanju W. Meng, Bojun Chen

## Abstract

Pre-mRNA splicing is essential for gene expression, yet the consequences of disrupting core spliceosomal factors during animal development remain poorly understood. Here, we investigated the *in vivo* function of the conserved spliceosomal kinase PRPF-4 in *C. elegans* using endogenous reporter analysis, conditional protein depletion, and transcriptome-wide analysis of alternative splicing and gene expression. We found that PRPF-4 is broadly expressed throughout development and is continuously required for postembryonic development and normal function of multiple tissues, including the pharynx, nervous system, and germline. Acute PRPF-4 depletion rapidly disrupts alternative splicing across thousands of transcripts, with exon skipping representing the predominant class of affected events. In addition, global PRPF-4 depletion causes widespread changes in gene expression, including increased expression of spliceosomal components and decreased expression of genes involved in ciliary function and ion transport, and neuron-specific depletion compromises amphid sensory cilia function. These findings establish PRPF-4 as an essential regulator of pre-mRNA splicing and demonstrate the widespread consequences of its depletion on gene expression, neuronal function, and animal development.

**Author Summary:** Pre-mRNA splicing, a process in which newly synthesized RNA molecules are cut and joined to produce mature RNA molecules, is essential for life. This study examines a conserved splicing protein called PRPF-4 in the nematode *C. elegans*. The researchers show that PRPF-4 is required throughout postembryonic development and for the normal function of multiple tissues. Depleting PRPF-4 causes numerous splicing errors and changes the expression of many genes, including those involved in cilia and ion transport. Neuronal depletion of PRPF-4 also impairs sensory cilia function. These findings establish PRPF-4 as an essential regulator of RNA processing during animal development.

## Introduction

Pre-mRNA splicing is a fundamental step in eukaryotic gene expression, which involves the excision of the non-coding sequences (introns) and the ligation of the coding regions (exons) to produce a mature messenger RNA (mRNA). This process is catalyzed by the spliceosome, a large, highly dynamic ribonucleoprotein complex that assembles from five small nuclear ribonucleoproteins (snRNPs; U1, U2, U4, U5 and U6) and many associated proteins (Will and Luhrmann 2011; Matera and Wang 2014). Beyond its universal role in ‘constitutive’ splicing, the use of different splice sites within a gene to generate different mRNA isoforms, known as ‘alternative splicing’, is the largest source of transcriptomic complexity in eukaryotic organisms (Nilsen and Graveley 2010; Baralle and Giudice 2017). In metazoans, alternative splicing is both widespread and fundamental to development, cell differentiation, and tissue-specific gene expression (Pan *et al*. 2008; Hyung *et al*. 2018; Koterniak *et al*. 2020; Mazin *et al*. 2021). Consistent with its central role in the regulation of gene expression, defects in pre-mRNA splicing are linked to a wide range of human diseases (Yoshida and ogawa 2014; Scotti and swanson 2016).

PRP4K (pre-mRNA processing factor 4, also known as PRPF4B) is an evolutionarily conserved serine/threonine kinase and a component of the U4/U6-U5 tri-snRNP of the pre-mRNA splicing machinery (Dellaire *et al*. 2002). In fission yeast and mammalian cells, PRP4K was shown to phosphorylate multiple spliceosomal components and play roles in spliceosome assembly and activation (Schwelnus *et al*. 2001; Lutzelberger *et al*. 2010; Schneider *et al*. 2010; Eckert *et al*. 2016). In human cells, PRP4K phosphorylates U5 snRNP proteins PRPF6 and PRPF31, thereby stabilizing the U4/U6-U5 tri-snRNP in the spliceosome and promoting the formation of the activated B complex (Schneider *et al*. 2010). More recently, PRP4K was shown to function in a highly evolutionarily conserved splicing pathway, which regulates autophagy in human cells and amoeba (Mathavarajah *et al*. 2025). In addition to its well-established function in pre-mRNA splicing, PRP4K has also been reported to be involved in cellular processes related to tumor suppression (Habib *et al*. 2022). Consistent with its essential function, siRNA-mediated depletion of the PRP4K homolog, PRPF-4, in the nematode *C. elegans* was shown to cause embryonic lethality (Dellaire *et al*. 2002), and in *Drosophila*, Prp4k has been associated with lethality in a P-element insertion mutant (Cho *et al*. 2018) and following RNAi-mediated depletion that also disrupts neural stem cell development (Neumuller *et al*. 2011). Thus, loss of PRP4K is generally not tolerated across metazoans. Despite these findings, the *in vivo* roles of PRP4K, such as its global effects on the transcriptome, remain undefined yet.

Here, we investigated the function of PRPF-4 in *C. elegans* using genetic, molecular, and transcriptomic approaches. We defined its expression pattern and examined the tissue requirements for PRPF-4 function through conditional depletion. We found that acute depletion of PRPF-4 caused widespread disruption of alternative splicing across all major classes of splicing events, as well as coordinated transcriptional alterations, such as increased expression of genes encoding spliceosomal proteins and decreased expression of genes involved in ciliary function and ion transport. These observations provide insight into the role of a conserved core spliceosomal kinase in maintaining RNA processing, gene expression, and normal development in metazoans.

## Materials and Methods

### *C. elegans* strains and culture conditions

All *C. elegans* strains were cultured on nematode growth medium (NGM) plates seeded with *Escherichia coli* OP50 at 20°C. The following strains were generated in this study: BJC161: *Phx7895(prpf-4a::SL2::mNeonGreen)*. BJC162: *Phx7876(prpf-4a::degron)*. BJC204: *Phx7876(prpf-4a::degron); ieSi38 [sun-1p::TIR1::mRuby::sun-1 3’UTR + Cbr-unc-119(+)]*. BJC205: *Phx7876(prpf-4a::degron); ieSi57 [eft-3p::TIR1::mRuby::unc-54 3’UTR + Cbr-unc-119(+)]*. BJC206: *Phx7876(prpf-4a::degron); reSi7 [rgef-1p::TIR1::F2A::mTagBFP2::AID::NLS::tbb-2 3’UTR]*. BJC262: *Phx7876(prpf-4a::degron); ieSi60 [myo-2p::TIR1::mRuby::unc-54 3’UTR + Cbr-unc-119(+)]*. The *Phx7876* and *Phx7895* alleles were generated using CRISPR/Cas9 genome editing in the N2 (Bristol) background by SunyBiotech. In *Phx7895*, an *SL2* trans-splicing acceptor followed by the coding sequence of a bright GFP variant, mNeonGreen, was inserted directly after the stop codon of the PRPF-4a isoform. In *Phx7876*, a degron tag is inserted at the C-terminus of the PRPF-4a isoform, immediately before the stop codon in the final exon. The degron tag sequence corresponds to the minimal auxin-inducible degron (AID) derived from *Arabidopsis thaliana* IAA17 (Nishimura *et al*. 2009), and was synthesized *de novo*. Both alleles showed no detectable phenotypes, suggesting that neither the SL2::mNeonGreen insertion nor the degron tag substantially disrupts PRPF-4 function. Strains carrying tissue-specific TIR1 transgenes for the AID experiments were obtained from the *Caenorhabditis Genetics Center* (CGC) and have been previously described (Zhang *et al*. 2015).

### PRPF-4 expression analysis and fluorescence microscopy

Worms expressing *prpf-4a::SL2::mNeonGreen* were mounted on 2% agarose pads in M9 buffer containing 10 mM sodium azide and imaged using a Leica THUNDER Imager 3D Tissue fluorescence microscope equipped with a Leica K8 digital camera. mNeonGreen fluorescence was examined in adult animals and at the L1, L2, L3, and L4 larval stages to characterize the spatial and developmental expression pattern of PRPF-4. Corresponding differential interference contrast (DIC) images were acquired for anatomical reference.

### Analysis of locomotion speed

Locomotion speed was analyzed using an automated worm-tracking and analysis system (Wang and Wang 2013). Briefly, a single young adult hermaphrodite was placed on an NGM plate without food and allowed to recover for approximately 30 s after transfer. Images of the worm were then acquired at 15 frames per second (fps) for 30 s using a digital camera (DMK37BUX273, The Imaging Source) mounted on a stereomicroscope (SMZ800, Nikon). The worm was continuously maintained in the center of the field of view using a motorized microscope stage (OptiScan ES111, Prior Scientific). Both the camera and the motorized stage were controlled by a custom-written program in MATLAB (The MathWorks). Images were analyzed using custom worm-tracking software written in MATLAB to quantify locomotion speed (Wang and Wang 2013).

### Real-time quantitative PCR (RT-qPCR)

Total RNA was extracted from mixed-stage worms using TRIzol Reagent (Invitrogen, 15596026) and treated with TURBO DNase (Ambion, AM1907). One microgram of total RNA was reverse transcribed with M-MuLV reverse transcriptase (New England BioLabs, M0253S) and an oligo(dT)16 primer. qPCR was performed in a 20 μl reation volume using iTaq Universal SYBR Green Supermix (Bio-Rad, 1725121) on a CFX Opus 96 Real-Time PCR System (Bio-Rad). The amplification conditions were as follows: 3 min at 95°C for initial polymerase activation, followed by 40 cycles of 10 s at 95°C for denaturation and 30 s at 62°C for annealing and extension. Relative mRNA levels were calculated using the comparative Ct (2L^ΔΔCt^) method and expressed as fold changes relative to the control. qPCR was performed in technical triplicates, and each biological sample was independently normalized to the endogenous reference gene *act-1*. The primers used for *prpf-4* were 5′-TGATGGGCATAGTGGAGCAG (forward) and 5′-TGCATCGGCATACATTACCAA (reverse). The primers used for *act-1* were 5′-GCCCAATCCAAGAGAGGTATCC (forward) and 5′-TGAGGAGGACTGGGTGCTCT (reverse).

### Dye-filling assays

Synchronized L3-stage BJC206 worms were transferred to NGM plates supplemented with 1 mM IAA (indole-3-acetic acid; ThermoScientific, A10556.14). Control animals were transferred to NGM plates without IAA. When the worms reached young adulthood, they were washed off the plates with M9 buffer and transferred to 1.5-mL microcentrifuge tubes. DiI stock solution (2.5 mg/mL in DMSO; Thermo Fisher Scientific, D3911) was added to the worm suspension at a 1:1,000 dilution. The worms were incubated with gentle shaking for 30 min at room temperature. Subsequently, the worms were briefly centrifuged and transferred to fresh NGM plates, and were allowed to recover and excrete excess dye for approximately 1 h before imaging. Worms were mounted on 2% agarose pads in M9 buffer containing 10 mM sodium azide and imaged using the Leica imaging system described above. Fluorescence intensity was quantified using Leica LAS X software. For each worm, the region of interest (ROI) encompassing the amphid cilia and their associated sensory neuronal cell bodies in the head was selected, and the mean fluorescence intensity within this region was measured. Background fluorescence was measured from an adjacent region without specific DiI fluorescence and subtracted from the mean fluorescence intensity of the ROI. Background-corrected mean fluorescence intensity was used for subsequent comparisons between control and PRPF-4-depleted worms.

### Measurement of worm body volume

Day 2 adult worms were transferred to fresh NGM plates without food, and images were acquired using an Olympus SZX16 stereomicroscope equipped with an Allied Vision Alvium 1800 digital camera. Images were first processed using Adobe Photoshop (CS5, version 12.0.3), and worm body volume was then automatically measured using the WormSizer plugin (version 1.2.5) for Fiji/ImageJ (version 1.54t), as described by Moore et al. (2013) (Moore *et al*. 2013).

### Analysis of reproductive phenotypes

To assess reproductive capacity over time, individual L4-stage hermaphrodites were placed on fresh NGM plates (day 1) and transferred to fresh plates daily until the cessation of egg-laying. Worms were maintained at 20°C and the number of progeny produced on each day was counted two days later. The total number of progeny (brood size) was calculated as the sum of all progeny produced by an individual worm throughout its reproductive period. Reproductive lifespan was defined as the number of days from the L4 larval stage until the production of the last progeny. Animals were considered to have ceased reproduction when no new progeny were observed for two consecutive days. The reproductive lifespan (in days) was recorded for individual animals of both genotypes.

### Auxin-inducible degradation (AID) experiments

For PRPF-4 depletion using AID, worms at different larval stages were transferred to NGM plates supplemented with 1 mM IAA (indole-3-acetic acid; ThermoScientific, A10556.14). Control animals were transferred to plates without IAA. For phenotypic analyses, worms were treated continuously for 72 hours at 20°C, and phenotypes were scored and representative images were acquired using an Olympus SZX16 stereo microscope equipped with an AmScope MU300 digital camera. IAA-containing plates were prepared fresh daily and protected from light.

### RNA isolation, library preparation and sequencing

For RNA-seq experiments, synchronized L4-stage BJC205 worms were treated with 1 mM IAA or mock-treated (no IAA) for 2 hours at 20°C on NGM plates seeded with *E. coli* OP50. Worms were then collected by washing three times with pre-cooled M9 buffer at 4°C. Three biological replicates were collected for each condition, with each replicate consisting of approximately 2,000 - 3,000 worms. Total RNA was extracted using TRIzol reagent (Invitrogen, 15596026) according to the manufacturer’s instructions. Directional sequencing libraries were prepared using the VAHTS Universal V10 RNA-seq Library Prep Kit for Illumina (Vazyme, NR606-01) according to the manufacturer’s instructions. Briefly, mRNA was purified from total RNA using poly-T oligo-attached magnetic beads. Following fragmentation, first-strand cDNA was synthesized using random hexamer primers, followed by second-strand cDNA synthesis using dUTP in place of dTTP. Libraries were prepared using end repair, A-tailing, adapter ligation, size selection, USER enzyme digestion, amplification and purification. Libraries were quantified and checked for size distributions by Qubit fluorometer, quantitative real-time PCR, and Bioanalyser. After quality control, libraries were pooled according to the effective concentration and the expected sequencing depth, and sequenced by Illumina platform to generate 150-bp paired-end reads. Sequencing was performed by Novogene, Inc.

### Sequencing data processing

Raw sequencing data in FASTQ format were first processed using trim galore (v0.6.10) (FELIX KRUEGER 2023) to remove adapter-containing reads, reads containing poly-N and low-quality reads. All downstream analyses were performed using the high-quality clean reads. Clean paired-end reads were aligned to the reference genome (WBcel235) using STAR (v2.7.11b) (Dobin *et al*. 2013). Gene expression levels were quantified using featureCounts (v2.1.1) (Liao *et al*. 2014) by counting reads mapped to annotated genes.

### Alternative splicing analysis

Alternative splicing (AS) events were identified and quantified using rMATS (v4.3.0) (Shen *et al*. 2012). Five major AS event types were analyzed: skipped exon (SE), alternative 5′ splice site (A5SS), alternative 3′ splice site (A3SS), mutually exclusive exon (MXE), and retained intron (RI). AS events with a false discovery rate (FDR) < 0.05 and absolute IncLevelDifference > 0.1 were considered statistically significant. Sashimi plots were generated by grouping replicates using rmats2sashimiplot (v4.0.0) (https://github.com/Xinglab/rmats2sashimiplot/).

### Differential expression analysis

Differential expression analysis between two conditions was performed using the DESeq2 R package (v1.46.0). The resulting *p*-values were adjusted using the Benjamini and Hochberg method to control the error discovery rate. Genes with an adjusted *p*-value (padj) ≤ 0.05 and an absolute log2(fold change) ≥ 1 were considered significantly differentially expressed.

### Gene Ontology and KEGG enrichment analysis

Gene Ontology (GO) enrichment analysis was performed using the clusterProfiler package v4.14.6 in R. GO terms with padj < 0.05 were considered significantly enriched. Kyoto Encyclopedia of Genes and Genomes (KEGG) pathway enrichment analysis was also performed using clusterProfiler with the *C. elegans* KEGG annotation database (org.Ce.eg.db, v3.22.0). Pathways with padj < 0.05 were considered significantly enriched.

### Gene network analysis

Functional association networks were generated using GeneMANIA (version 3.6.0) (Warde-Farley *et al*. 2010) via its web interface (https://genemania.org). Network weighting was performed using the automatically selected weighting method, which integrates multiple evidence sources, including co-expression, predicted functional relationships, physical protein–protein interactions, genetic interactions, and shared protein domains. To restrict the network to the query genes and their direct associations, both max resultant genes and max resultant attributes were set to 0. The resulting composite network was visualized with edges color-coded according to evidence category and nodes labeled with gene names. The percentage contribution of each evidence type to the final network is reported in the figure legend.

## Results

### PRPF-4 is broadly expressed throughout development in *C. elegans*

To determine the native expression pattern of *prpf-4*, we generated a transcriptional reporter strain through the CRISPR/Cas9 approach. An *SL2* trans-splicing acceptor followed by the coding sequence of a bright GFP variant, *mNeonGreen*, was inserted directly after the stop codon of the major splicing variant *prpf-4a* (**Fig. 1a**). This bicistronic design allows the native *prpf-4* promoter to drive transcription of both *prpf-4* and *mNeonGreen*. Because the open reading frames of *prpf-4* and *mNeonGreen* are separated by the *SL2* sequence, the two proteins are translated independently, and the mNeonGreen fluorescence is representative of cells expressing native *prpf-4*. The reporter strain was indistinguishable from wild type worms morphologically and behaviorally, indicating this insertion does not disrupt *prpf-4* expression or function.

**Fig. 1.**
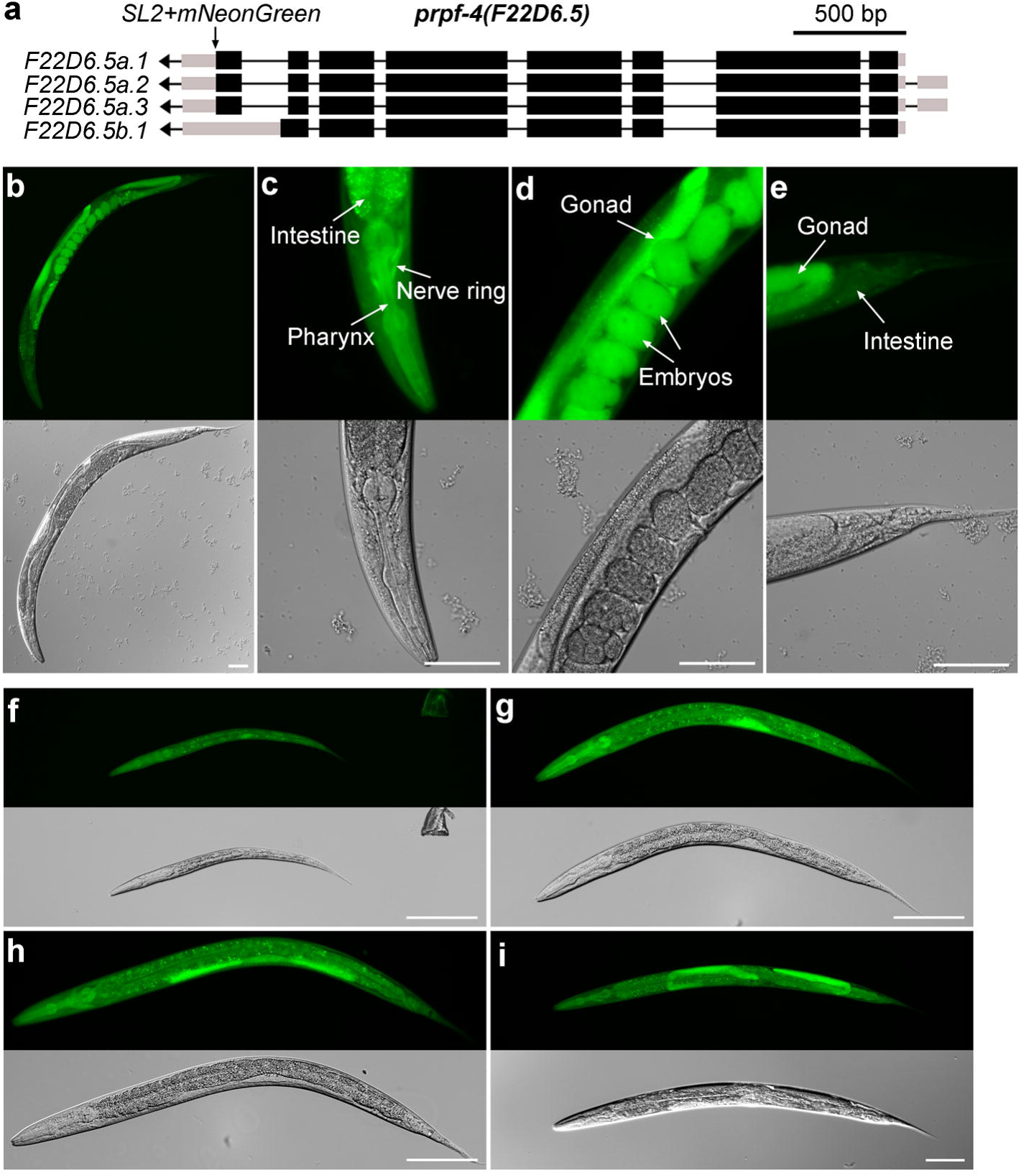
PRPF-4 is broadly expressed throughout development. a) Schematic of the *prpf-4* locus showing the PRPF-4 isoforms and the position of the *SL2::mNeonGreen* cassette inserted downstream of the *prpf-4a* coding sequence. Black boxes and lines represent exons and introns, respectively, and gray boxes represent untranslated regions. b - e) Representative mNeonGreen fluorescence images of adult *Phx7895* (*prpf-4a::SL2::mNeonGreen*) animals, showing broad PRPF-4 expression throughout the animal, including the intestine, pharynx, nerve ring, and gonad. Embryos within the gonad also showed strong fluorescence. f - i) Representative mNeonGreen fluorescence images of *Phx7895* animals at the L1 (f), L2 (g), L3 (h), and L4 (i) larval stages, respectively, demonstrating PRPF-4 expression throughout postembryonic development. Corresponding DIC images are shown below the fluorescence images. Scale bars = 50 µm.

Fluorescence microscopy of the reporter strain revealed that *prpf-4* is expressed throughout *C. elegans* development. In adult animals, the reporter protein was widely expressed across tissues, with high expression levels observed in the nervous system, pharynx, and the gonadal lumen (**Fig. 1b -e**). A very strong signal was also observed in early-stage embryos contained within the mother (**Fig. 1b** and **d**).

Since zygotic transcription is very limited and maternal products still dominate at these early embryonic stages (2-to 4-cell stages)(Robertson and Lin 2015), this embryonic signal likely reflects maternally deposited mNeonGreen protein synthesized in the germline. These observations suggest that the *prpf-4* promoter is also active in adult hermaphrodite reproductive tissue, and that the reporter protein is transmitted to the next generation through the maternal cytoplasm.

We also examined the reporter expression at different larval stages and found its expression is maintained throughout postembryonic development. Strong mNeonGreen fluorescence was readily detected in L1, L2, L3, and L4 larvae (**Fig. 1f - i**), with signal distributed throughout the body at each stage. No obvious stage-specific restriction or loss of expression was observed, indicating that *prpf-4* is continuously expressed during larval development as well as in adulthood. The widespread, constitutive distribution of the reporter throughout development is consistent with the expected ubiquitous requirement of PRPF-4 as a core component of the spliceosome.

### PRPF-4 is required throughout larval development

To investigate the *in vivo* function of PRPF-4, we attempted to generate a *prpf-4* null mutant using the CRISPR/Cas9 approach. However, no viable homozygous knockout animals were recovered, and heterozygous animals produced dead embryos, indicating that *prpf-4* is essential for embryonic viability. This observation is consistent with the essential role of PRP4K homologs in other species(Alahari *et al*. 1993; Habib *et al*. 2022). To bypass the embryonic lethality and investigate the tissue-specific requirements for PRPF-4, we next employed the auxin-inducible degradation (AID) system to achieve temporal and spatial depletion of PRPF-4. We tagged the endogenous *prpf-4* locus at its C-terminus with a degron sequence and crossed this allele into strains expressing the TIR1 E3 ligase in various tissue-specific backgrounds. The *prpf-4::degron* worms ware indistinguishable from *wild type* worms in *prpf-4* splicing patterns, mRNA levels, locomotion speed, body volume, and reproductive phenotypes (**Supplementary Fig. 1**), indicating that the C-terminal degron-tagging does not interfere with normal PRPF-4 function.

To determine the temporal requirement for PRPF-4, we depleted endogenous PRPF-4 at different stages of postembryonic development. Synchronized L1-, L2-, L3-, or L4-stage *prpf-4::degron* animals expressing TIR1 in all somatic cells were grown on NGM plates containing either 0 mM or 1 mM IAA (indole-3-acetic acid, IAA). In the absence of IAA, worms developed normally through all larval stages (**Fig. 2a**). In contrast, exposure to IAA caused developmental arrest regardless of the larval stage at which treatment was initiated, and some animals subsequently died during continued treatment (**Fig. 2a**). These results demonstrate that PRPF-4 is required throughout postembryonic development and is essential beyond a single developmental window.

**Fig. 2.**
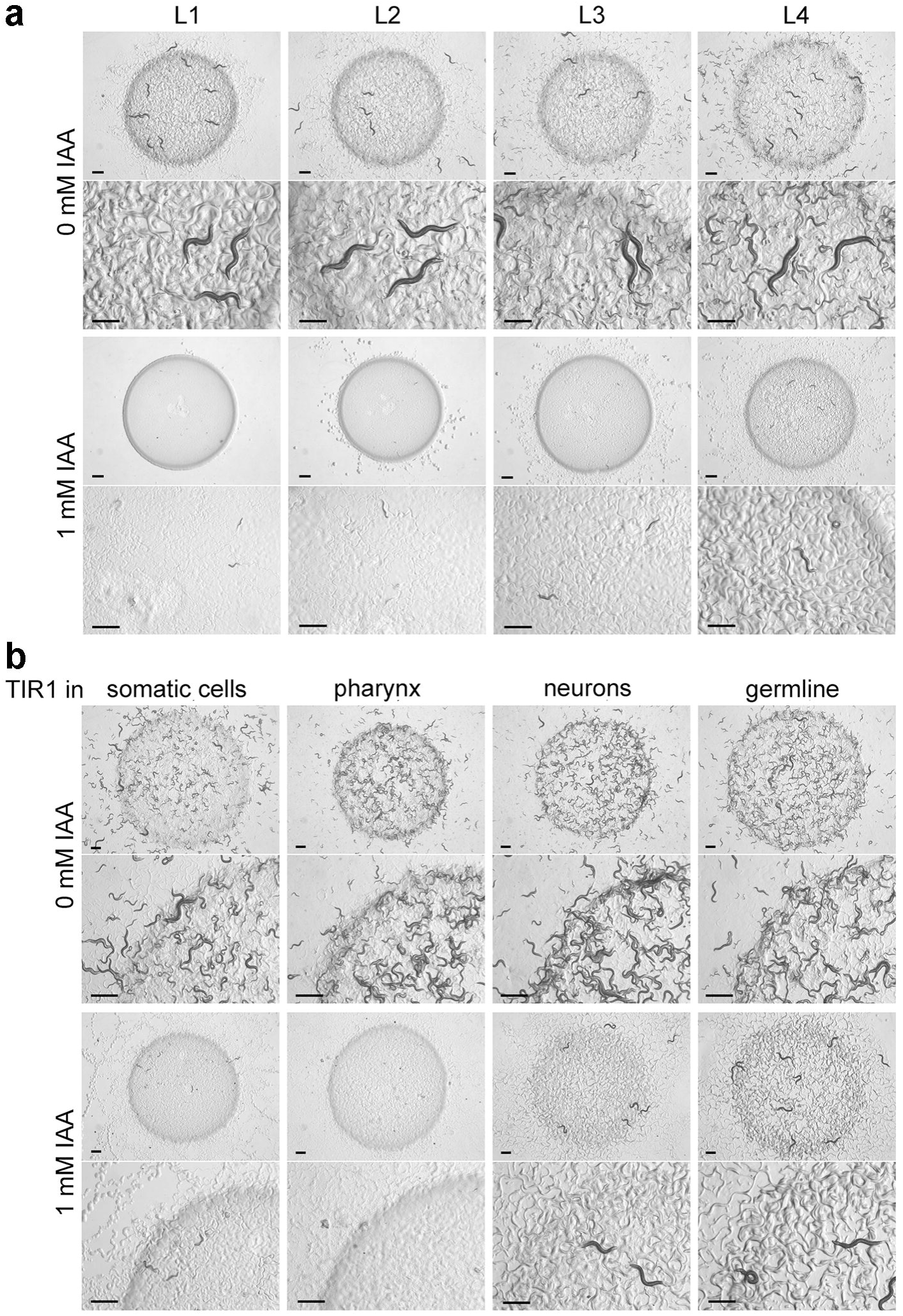
PRPF-4 is essential in multiple cell types during *C. elegans* development. a) Effect of PRPF-4 depletion in all somatic cells during development. Representative images are shown for worms expressing *prpf-4::degron* and TIR1 in all somatic cells treated with or without IAA. Ten worms at the L1 - L4 stages were placed on NGM plates containing 1 mM IAA or control plates without IAA, and images were taken after three days of incubation at 20 °C. Untreated animals developed normally, whereas auxin-induced degradation of PRPF-4 at any larval stage resulted in growth arrest, with some animals died during continued treatment. The strain used was *Phx7876;ieSi57*. In *Phx7876*, PRPF-4 is tagged with a degron at the C-terminus. In *ieSi57*, TIR1 is expressed in all somatic cells. b) Effect of PRPF-4 depletion in different cell types. Representative images are shown for worms expressing *prpf-4::degron* and TIR1 in specific tissues and treated with or without IAA. Ten L4-stage worms from each strain were placed on NGM plates containing 1 mM IAA or control plates without IAA, and images were taken after three days of incubation at 20 °C. Auxin-induced degradation of PRPF-4 in somatic cells, the pharynx, or neurons caused growth arrest, whereas germline-specific depletion produced sterile adults without an obvious developmental growth defect. The strains used were *Phx7876;ieSi57* for PRPF-4 depletion in all somatic cells, *Phx7876;ieSi60* for pharynx, *Phx7876;ieSi7* for neurons, and *Phx7876;ieSi38* for germline. Scale bars = 500 µm.

To define potential tissue-specific requirements for PRPF-4, we crossed *prpf-4::degron* into strains expressing TIR1 in different tissues and examined the effect of IAA treatment. We selected the pharynx, nervous system, and germline for tissue-specific depletion because these tissues have relatively high *prpf-4* expression (**Fig. 1**), and because previous studies have shown extensive alternative splicing and distinct tissue-specific splicing programs in these tissues (Ragle *et al*. 2015; Watabe *et al*. 2018; Koterniak *et al*. 2020; Weinreb *et al*. 2025). Depletion of PRPF-4 specifically in the pharynx caused developmental arrest and sometimes death, which is similar to the phenotype observed following depletion in all somatic cells (**Fig. 2b**). Depletion of PRPF-4 from all neurons caused a mild developmental delay, but the animals were able to reach adulthood and produced a small number of embryos. However, these embryos either failed to hatch or were arrested shortly after hatching (**Fig. 2b**), suggesting that neuronal PRPF-4 function is also required for normal development. In contrast, depletion of PRPF-4 in the germline did not prevent animals from developing into adults but resulted in complete sterility (**Fig. 2b**), indicating that PRPF-4 in the germline is dispensable for somatic development but essential for germline function.

Together, these findings demonstrate that *prpf-4* is an essential gene with broad requirements in different tissues. PRPF-4 activity in the pharynx and other somatic tissues is required for postembryonic growth and viability, whereas its activity in the germline is specifically required for fertility. Although neuronal depletion produced only mild defects in the treated animals, the resulting arrested phenotype of their progeny indicates that PRPF-4 function in the nervous system is also critical for normal development. These results are consistent with previous findings that siRNA-mediated depletion of PRPF-4 in *C. elegans* causes embryonic lethality (Dellaire *et al*. 2002), and with the essential role of PRP4K homologs in other species (Alahari *et al*. 1993; Cho *et al*. 2018; Kanno *et al*. 2018).

### Acute PRPF-4 depletion leads to widespread alternative splicing changes

To investigate the molecular function of PRPF-4 in splicing regulation *in vivo*, we performed RNA-seq on worms subjected to acute PRPF-4 depletion (2-hour IAA treatment) and on worms of the same strain maintained without IAA treatment. Differential alternative splicing (AS) analysis identified 18,650 AS events affecting a total of 6,269 genes. Using an FDR threshold of < 0.05 and absolute IncLevelDifference > 0.1, we identified 602 significant events with increased exon inclusion and 1,930 significant events with decreased exon inclusion relative to untreated controls (**Fig. 3a** and **Supplementary Table 1**). The distribution of IncLevelDifference values indicated that most significant events exhibited weak-to-moderate changes in exon inclusion, although a subset showed large changes in exon inclusion (**Fig. 3b**).

**Fig. 3.**
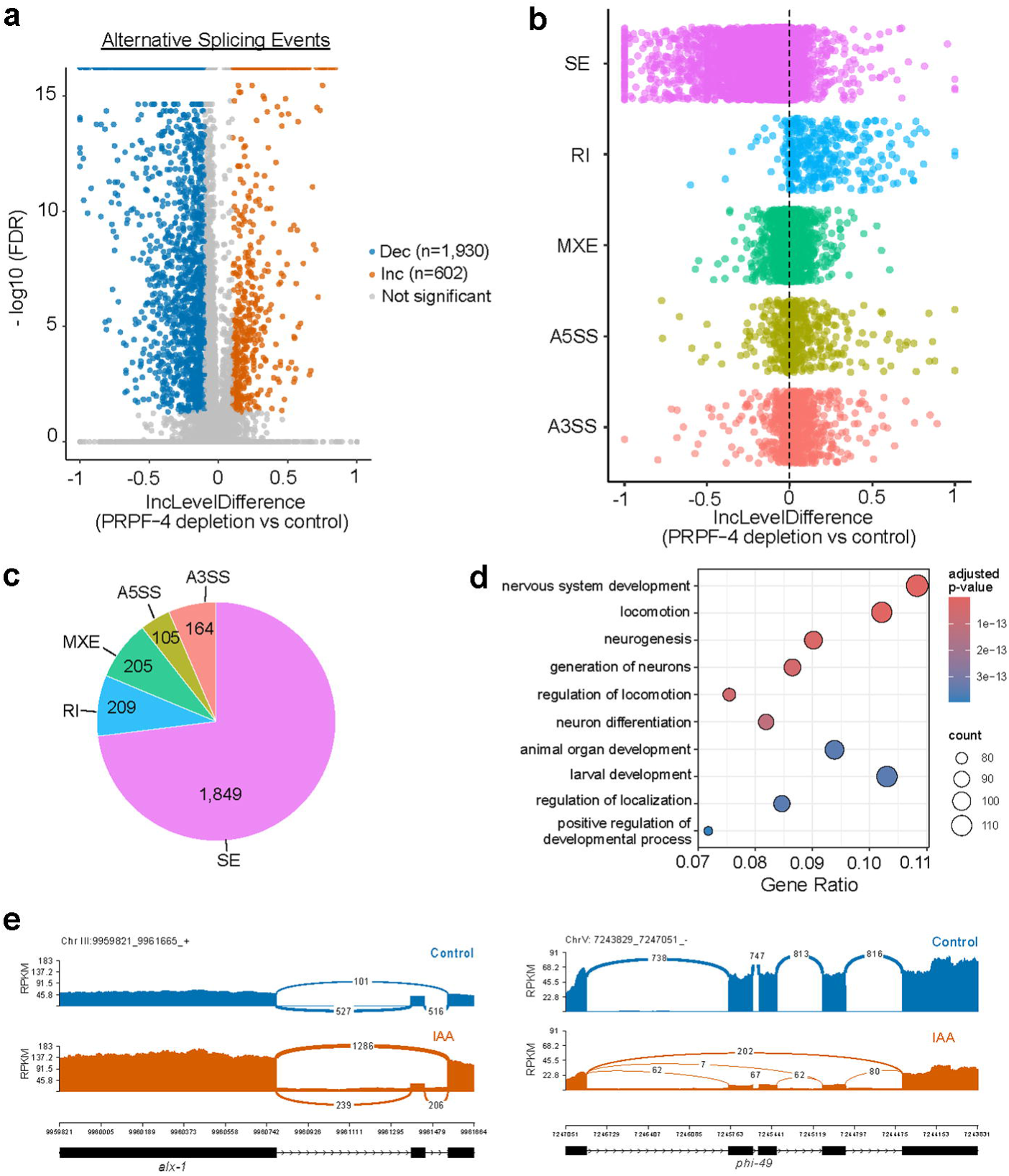
Acute depletion of PRPF-4 causes widespread alternative splicing changes. a) Volcano plot of alternative splicing events following acute depletion of PRPF-4. Each dot represents one alternative splicing event. The x-axis indicates the difference in exon inclusion levels (IncLevelDifference), and the y-axis indicates statistical significance expressed as −log10(FDR). Orange and blue dots represent significantly increased (IncLevelDifference > 0.1) and decreased (IncLevelDifference < 0.1) exon inclusion, respectively, while gray dots indicate events that did not reach statistical significance. A total of 1,930 events showed increased exon inclusion and 602 showed decreased exon inclusion (FDR < 0.05). b) Distribution of exon inclusion-level differences for each class of alternative splicing event. Each point represents an individual alternative splicing event, illustrating the range and frequency of splicing changes in each class of alternative splicing. c) Distribution of significant alternative splicing events among the five major event types: skipped exon (SE) events were the predominant class, accounting for 1,849 events, followed by retained intron (RI, 209), mutually exclusive exons (MXE, 205), alternative 3′ splice site (A3SS, 164), and alternative 5′ splice site (A5SS, 105). d) Dot plot showing top 10 enriched GO Biological Process terms among genes affected by alternative splicing following PRPF-4 depletion. Dot size represents the number of genes associated with each GO term, and color indicates the adjusted P value. Enriched terms were predominantly related to nervous system development, locomotion, and neurogenesis. e) Representative sashimi plots showing alternative splicing changes following PRPF-4 depletion. RNA-seq read coverage (RPKM) is shown for control and IAA-treated animals at the *alx-1* and *phi-49* loci. Junction reads are indicated by arc numbers. Gene models are shown below each coverage plot, with exons represented by black boxes and the direction of transcription indicated by arrows.

Among the five major classes of AS, skipped exons (SE) were the most prevalent, accounting for 1,849 events (73.0% of all significant AS events). The remaining four classes occurred at substantially lower frequencies: retained introns (RI), 209 events; mutually exclusive exons (MXE), 205 events; alternative 3′ splice sites (A3SS), 164 events; and alternative 5′ splice sites (A5SS), 105 events (**Fig. 3c**). Consistent with their high frequency, the distributions of inclusion-level differences showed that SE events exhibited the broadest range and largest magnitude of splicing changes among the five AS classes (**Fig. 3b**). These significant AS events corresponded to a nonredundant set of 1,586 unique genes, indicating that PRPF-4 broadly influences alternative splicing decisions. In addition, as shown in representative examples of altered splicing events in **Fig. 3e**, PRPF-4 may affect different types of splicing events in different ways.

To further examine the functional effects of these splicing changes, we performed Gene Ontology (GO) enrichment analysis of the 1,586 affected genes. Our analysis revealed enrichment for developmental and nervous system-related biological processes. The most significantly enriched biological processes included nervous system development, locomotion, neurogenesis, larval development, and animal organ development (**Fig. 3d**). Overall, our results suggest that acute loss of PRPF-4 results in a rapid disruption of exon choice in many different transcripts, with exon skipping as the major splicing error and a clear bias toward genes involved in development of the nervous system and organism developmental processes.

### Acute PRPF-4 depletion induces a compensatory transcriptional response

In addition to alternative splicing changes, we investigated global gene expression changes upon acute PRPF-4 depletion. Differential expression analysis identified 3,239 up-regulated genes and 3,009 down-regulated genes (**Fig. 4a** and **Supplementary Table 2&3**). Notably, GO enrichment analysis of the up-regulated genes revealed strong enrichment for RNA splicing (**Fig. 4b**). The two most significant enriched GO terms were RNA splicing and RNA splicing via transesterification reactions, with more than 50 known splicing-related genes significantly induced. Similar findings were observed in KEGG pathway analysis of up-regulated genes, in which the spliceosome was the most significantly enriched pathway (**Supplementary Fig. 2a**). Interestingly, most of the upregulated splicing-related genes did not show significant alternative splicing changes (**Supplementary Table 4**), suggesting their upregulation is predominantly associated with transcriptional rather than alternative splicing changes. In addition to RNA splicing and mRNA processing, several other highly enriched GO categories were related to host defense response (e.g. defense response to other organisms, response to biotic stimulus) (**Fig. 4b**). This observation is consistent with previous findings linking RNA splicing factors to immune and cellular stress responses in *C. elegans (Kew et al. 2020; Xia et al. 2022)*.

**Fig. 4.**
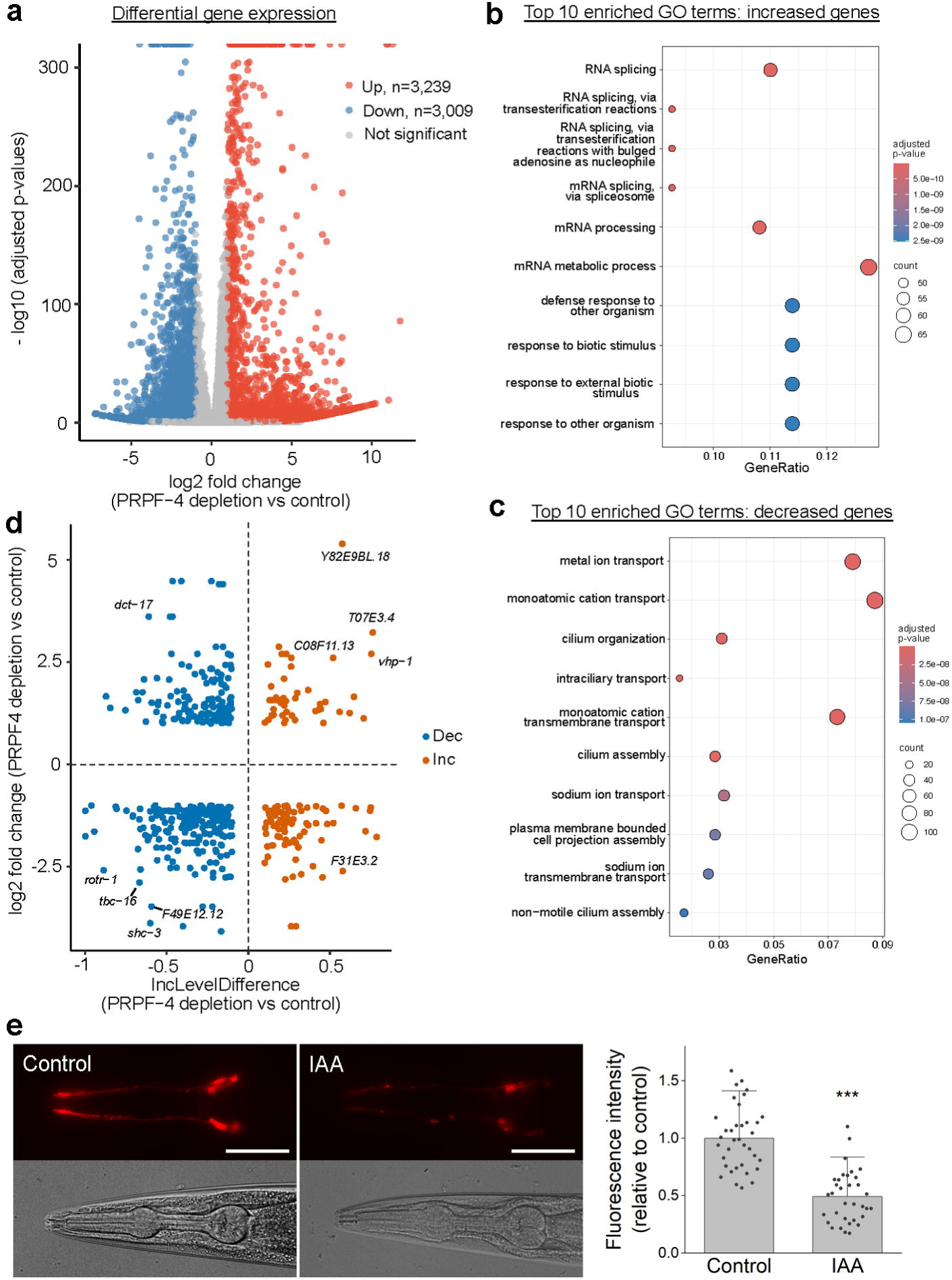
Differential gene expression upon PRPF-4 depletion reveals a compensatory transcriptional response. a) Volcano plot showing differentially expressed genes in PRPF-4 depleted worms compared to control. Significantly upregulated genes (n=3,239) are shown in red, and downregulated genes (n=3,009) are shown in blue, with the threshold set at |log2(fold change)| > 1 and adjusted *p*-value < 0.05. b) Dot plots showing top 10 enriched GO Biological Process terms among up-regulated genes following PRPF-4 depletion. Up-regulated genes are predominantly enriched for RNA splicing. c) Dot plots showing top 10 enriched GO Biological Process terms among down-regulated genes following PRPF-4 depletion. Down-regulated genes are enriched for ion transport and ciliary organization. In b) and c), the dot size represents the number of genes associated with each GO term, and the color indicates the adjusted P value. d) Relationship between differentially expressed genes and genes showing significant changes in alternative splicing following acute PRPF-4 depletion. Each point represents an alternative splicing event. Blue and orange points indicate genes with decreased (Dec) and increased (Inc) exon inclusion, respectively. The number of alternative splicing (AS) events and genes are indicated in each quadrant. Selected genes with large changes in exon inclusion and gene expression are labeled. e) Representative fluorescent and corresponding DIC images, and comparison of fluorescence intensity between control (n = 36) and PRPF-4-depleted worms (n = 34) treated with DiI. Data are shown as mean, and error bars indicate +1 SD. (*** *p* < 0.001, Welch’s two-sample t-test.). Scale bars = 50 µm.

In contrast, GO enrichment analysis of the down-regulated genes revealed a distinct set of functional categories, most notably related to ion transport and ciliary processes (**Fig. 4c**). Metal ion transport was the most significantly enriched term in the down-regulated genes followed by monoatomic cation transport, ciliary organization, and intraciliary transport. Enrichment was also significant for cilium assembly, sodium ion transport, plasma membrane-bounded cell projection assembly, and non-motile cilium assembly. Analysis of the down-regulated genes using KEGG pathway analysis further revealed significant enrichment for genes involved in axon regeneration, glycerophospholipid metabolism, IgSF CAM signaling, and the phosphatidylinositol signaling system (**Supplementary Fig. 2b**). Together, these results indicate that acute loss of PRPF-4 function is associated with coordinated down-regulation of genes involved in ciliary function and membrane ion homeostasis, as well as membrane lipid and phosphoinositide-related pathways.

To understand the functional relationships among the down-regulated cilia-associated genes identified via GO analysis (**Supplementary Table 4**), we constructed a functional association network using GeneMANIA (Warde-Farley *et al*. 2010). As shown in **Supplementary Fig. 3**, the network is dominated by co-expression relationships, which account for 59.30% of the interactions, suggesting that these genes are transcriptionally regulated in a coordinated manner. The analysis also revealed a dense sub-network of predicted association (29.81%) centered on the BBSome complex components (e.g., *bbs-1, bbs-4, bbs-5, bbs-8,* and *bbs-9*), BBSome-associated trafficking factors (e.g., *arl-6*), and intraflagellar transport (IFT) components (e.g., *osm-5*, *osm-12*, *che-11*, and *ifta-1*). This cluster suggests the functional convergence of these genes on sensory cilium organization and assembly. Together, these analyses indicate that the cilia-associated genes affected by PRPF-4 depletion form a functionally cohesive and highly interconnected network.

We also constructed an association network among ion transport-related genes identified from the GO analysis (**Supplementary Table 4**). In contrast to the cilia-related network, this network is heavily driven by shared protein domains (45.25%) (**Supplementary Fig. 4**), suggesting that many of these genes encode proteins belonging to the same structural families. The network reveals two notable clusters, one comprising numerous *twk* genes encoding TWK-type potassium channels, and the other dominated by *del* and *deg* genes encoding DEG/ENaC sodium channels. Co-expression also accounts for a major portion of the associations (43.07%), suggesting that many of these ion transport-related genes are coordinately expressed. The representation of these distinct ion channel families (TWK and DEG/ENaC) within a highly interconnected network suggests that PRPF-4 depletion affects multiple components of ion homeostasis and membrane transport.

To determine the functional consequences of PRPF-4 depletion, particularly its potential effects on sensory cilia, we performed a DiI dye-filling assay with the BJC206 strain that allows neuron-specific PRPF-4 depletion. DiI dye-filling assay is a commonly used method for assessing the structural and functional defects in ciliated sensory neurons in *C. elegans (Andrews et al. 2025)*. IAA treatment was initiated at the L3 larval stage, and DiI staining was performed when the animals reached the young adult stage to assess the effects of PRPF-4 depletion during late larval development and early adulthood.

Consistent with the GO analysis showing enrichment of genes associated with ciliary functions among those affected by PRPF-4 depletion (**Fig. 4c**), we observed significantly reduced DiI fluorescence in the amphid sensory neurons of animals with neuron-specific PRPF-4 depletion compared with control animals (**Fig. 4e**), suggesting that neuronal depletion of PRPF-4 compromises sensory cilia integrity or dye accessibility.

Finally, we examined the relationship between transcriptional and post-transcriptional regulation by comparing differentially expressed genes with those showing significant changes in alternative splicing. A total of 332 alternatively spliced genes (492 AS events) also exhibited differential gene expression (**Fig. 4d** and **Supplementary Table 5**). Among these, 38 genes showed increased transcript levels accompanied by increased exon inclusion, whereas 108 genes showed increased transcript levels with decreased exon inclusion. Conversely, 65 genes displayed increased exon inclusion despite decreased transcript levels, while 154 genes showed decreased exon inclusion together with decreased transcript levels. We also examined the relationship between alternative splicing event types and changes in transcript abundance by classifying differentially expressed genes according to their associated alternative splicing event types (**Supplementary Table 5**). In four of the five event categories, a greater number of genes showed decreased expression than increased expression, including A3SS (19 vs. 10), A5SS (19 vs. 5), RI (28 vs. 15), and SE (152 vs. 104). In contrast, MXE events showed an opposite trend, with more genes exhibiting increased expression than decreased expression (21 vs. 14). Together, these results indicate that, although acute PRPF-4 depletion induces widespread changes in both gene expression and alternative splicing, the direction of these changes is not uniformly concordant, suggesting that transcriptional and splicing responses are only partially coupled.

Collectively, these data reveal two transcriptional programs triggered by acute depletion of PRPF-4: up-regulation of genes encoding proteins involved in RNA splicing and defense response, and down-regulation of genes encoding proteins involved in ciliary function and ion transport. These concurrent programs of gene regulation, in addition to widespread changes in alternative splicing, may contribute to the observed growth arrest and developmental defects.

## Discussion

In this study, we combined tissue-specific auxin-inducible degradation (AID), RNA-seq, alternative splicing and differential gene expression analysis to investigate the *in vivo* function of the conserved splicing kinase PRPF-4 in *C. elegans*. We demonstrate that PRPF-4 is required for viability and development, with a particularly pronounced requirement in the pharynx, nervous system, and germline, and that acute PRPF-4 depletion causes widespread changes across all five major classes of alternative splicing events, along with a dramatic transcriptome response characterized by increased expression of spliceosome components and decreased expression of genes involved in cilium biogenesis and ion transport. Importantly, our DiI dye-filling experiments provide functional evidence supporting these transcriptomic findings, as neuron-specific depletion of PRPF-4 significantly reduced dye filling in amphid sensory neurons, suggesting impaired sensory cilia integrity or accessibility. Collectively, these results establish PRPF-4 as a core regulator of RNA processing and reveal broader transcriptional consequences of impaired spliceosome function.

The broad expression of PRPF-4 is consistent with the fact that PRPF-4 is a core component of the U4/U6-U5 tri-snRNP complex (Dellaire *et al*. 2002), in which it promotes spliceosome assembly by phosphorylating the components and facilitating formation of the spliceosomal B complex (Schneider *et al*. 2010). The continuous requirement for PRPF-4 throughout larval development therefore suggests that its function is not restricted to a specific developmental transition but is required to sustain gene expression during ongoing growth. Consistent with its broad expression, we found that PRPF-4 is essential in all tissues examined, including the pharynx, neurons, and germline. These observed phenotypes may reflect the general cellular requirement for PRPF-4 rather than tissue-specific functions. Future studies systematically depleting PRPF-4 in additional tissues will be needed to determine whether certain tissues are more sensitive to PRPF-4 loss than others, and whether there are distinct tissue-specific requirements for this core spliceosomal factor.

Previous studies have shown that PRP4K orthologs play important roles in pre-mRNA splicing in different organisms. In fission yeast, Prp4 kinase activity is required for the splicing of many introns, particularly those with weak exon1/5’ splice sites and weak branch sequences (Eckert *et al*. 2016). In *Fusarium graminearum*, FgPrp4 is important for spliceosomal B-complex activation and splicing efficiency, although it is not essential for intron splicing (Gao *et al*. 2016). In *Arabidopsis thaliana*, mutations in PRP4KA lead to widespread and partially overlapping perturbations in alternative splicing and a range of developmental defects (Kanno *et al*. 2018). Consistent with these findings, we observed widespread alternative splicing defects following PRPF-4 depletion. Interestingly, our results show that skipped-exon events predominated among the affected splicing events, suggesting that PRPF-4 may play a particularly important role in exon recognition and definition. Furthermore, acute AID-mediated depletion caused extensive splicing changes within just 2 hours, affecting 2,532 significant alternative splicing events in 1,586 genes. The rapid appearance of these splicing changes following acute PRPF-4 depletion argues that they are closely associated with the loss of PRPF-4 rather than being a potential secondary consequence of developmental defects.

Skipped-exon events accounted for approximately 73% of the significant alternative splicing events, indicating that exon-definition decisions are particularly sensitive to PRPF-4 depletion. PRPF-4 is unlikely to function as a conventional sequence-specific alternative splicing regulator; instead, its role in spliceosome assembly may influence the efficiency or fidelity with which individual exons are recognized. Because alternative splicing is controlled by coordinated interactions among core spliceosomal components, RNA-binding proteins, and cis-regulatory sequences (Wang and Burge 2008; Fu and Ares 2014), disruption of a core spliceosomal activity could disproportionately affect exons with intrinsically weaker or more highly regulated splice sites. The enrichment of developmental and nervous system-related genes among the affected transcripts further suggests that PRPF-4-dependent splicing may be particularly important for gene expression programs requiring extensive transcript diversification.

Previous studies of PRP4K orthologs focused primarily on splicing defects and phenotypic consequences (Eckert *et al*. 2016; Gao *et al*. 2016; Kanno *et al*. 2018). Our transcriptomic analysis revealed that, in addition to the widespread alternative splicing defects, PRPF-4 depletion also causes extensive changes in transcript abundance. More than 3,200 genes were up-regulated, with strong enrichment for genes involved in RNA splicing and spliceosome-associated functions. The increased expression of more than 50 known splicing-related genes suggests that loss of PRPF-4 triggers a compensatory transcriptional response. However, the mechanisms by which PRPF-4 depletion is sensed and this transcriptional program is activated remain unknown. Furthermore, whether the increased expression of other splicing factors can partially compensate for PRPF-4 loss, and whether this response is conserved in other organisms, remain to be determined.

An important finding from our AID experiments is that PRPF-4 is required not only during development but also for post-developmental viability. Depletion of PRPF-4 at any larval stage resulted in developmental arrest, and prolonged depletion led to death in some animals. This suggests that PRPF-4 is required continuously to sustain gene expression and maintain cellular homeostasis, rather than only at specific developmental transitions. This observation is consistent with the essential role of PRP4K in spliceosome assembly and activation (Schneider *et al*. 2010), and it implies that loss of PRP4K function in adult tissues could have substantial consequences for tissue maintenance and function.

The partial overlap between differentially expressed genes and alternatively spliced genes suggests that transcriptional and post-transcriptional responses are related but not tightly coupled. Only 332 genes were affected at both levels, and the direction of changes in transcript abundance did not consistently correspond to the direction of exon inclusion. These observations argue against a simple model in which differential expression is merely a consequence of altered splicing. Instead, PRPF-4 depletion appears to trigger coordinated but partially independent changes in both transcript processing and gene expression.

In contrast to the strong enrichment of RNA-processing functions among up-regulated genes, genes that were down-regulated after PRPF-4 depletion showed enrichment for ciliary processes, ion transport, regulation of membrane potential, and other membrane-related pathways. These changes may contribute to the developmental and neuronal phenotypes observed after depletion of this core splicing factor, since ion channels and other membrane-targeted proteins are subjected to extensive alternative splicing in neurons (Weinreb *et al*. 2025). Consistent with a possible disruption of ciliary function, neuron-specific depletion of PRPF-4 reduced DiI uptake by amphid sensory neurons L a phenotype commonly linked to defects in the structure or function of sensory cilia (Bacaj *et al*. 2008; Power *et al*. 2020; Power *et al*. 2024). Loss of ion transport and membrane potential regulators may also impair neuronal excitability and signal transduction. Together, our transcriptomic and functional analyses suggest that PRPF-4 contributes to the maintenance of sensory cilia function, potentially through its broad role in regulating RNA processing and gene expression.

Overall, our results demonstrate that PRPF-4 is a broadly expressed and essential regulator of pre-mRNA splicing in *C. elegans*. Acute depletion of PRPF-4 rapidly causes changes in alternative splicing, preferentially exon skipping, and results in a compensatory induction of splicing pathway genes and repression of genes involved in cilia and ion transport functions. These data suggest that acute loss of PRPF-4 primarily causes a defect in spliceosome function, followed by broader changes in gene expression patterns that may contribute to the developmental phenotypes. Future functional studies of individual PRPF-4-dependent splicing events will help understand how disruption of this essential spliceosomal kinase may affect development and neuronal function, and provide broader insight into the biological consequences of spliceosome dysfunction.

## Data availability

The RNA-seq data generated in this study have been deposited in the Gene Expression Omnibus (GEO) under accession number GSE344094 (to be released upon publication). All other data are available in the main text or supplementary materials. Strains are available upon request.

## Acknowledgements

This work was supported by National Institute of Health (R35GM139620 to B.C). Some strains were provided by the CGC, which is funded by NIH Office of Research Infrastructure Programs (P40 OD010440).

## Supporting Information

**Supplementary Fig. 1.** C-terminal degron tagging of PRPF-4 does not affect its mRNA expression or function. a) Schematic representation of the *prpf-4* locus showing *prpf-4* splicing variants and the position of *degron* insertion. Black boxes and lines represent exons and introns, respectively, and gray boxes represent untranslated regions. b) Sashimi plots of alternative splicing at the *prpf-4* gene locus comparing RNA-seq data of *wild type* and *prpf-4::degron* worms. RNA-seq data for *wild type* worms were obtained from NCBI GEO: GSE315272 (Niu *et al*. 2026). c) Comparison of *prpf-4* mRNA level between *wild type* and *prpf-4::degron* worms. RT-qPCR were performed using three biological replicates. d) Representative images and comparison of body volume between *wild type* (n = 45) and *prpf-4::degron* (n = 49) worms. Scale bars = 100 µm. e) Comparison of locomotion speed between *wild type* (n = 14) and *prpf-4::degron* (n = 16) worms. f) Daily progeny counts of individual *wild type* and *prpf-4::degron* worms throughout their reproductive lifespans. g) Comparison of total progeny count between *wild type* and *prpf-4::degron* worms. h) Comparison of reproductive lifespan between *wild type* and *prpf-4::degron* worms. 8 *wild type* and 7 *prpf-4::degron* worms were analyzed in f – h. Data in all bar graphs are shown as means, and error bars indicate +1 SD. Statistical significance was determined using Welch’s two-sample t-test.

**Supplementary Fig. 2.** KEGG pathway enrichment analysis of differentially expressed genes following PRPF-4 depletion. a) Dot plot showing top enriched KEGG pathways for up-regulated genes. b) Dot plot showing top enriched KEGG pathways for down-regulated genes. Dot size represents gene count, and color indicates adjusted p-value (FDR).

**Supplementary Fig. 3.** GeneMANIA interaction network of cilia-related genes. The network displays the associations among 38 cilia-related genes identified from GO term analysis of the PRPF-4 depletion vs. control dataset. Nodes represent individual genes, and edges represent five types of interaction evidence, as indicated by different colors. The central cluster highlights a highly connected group of genes involved in intraflagellar transport (IFT) and BBSome function.

**Supplementary Fig. 4.** GeneMANIA interaction network of ion transport-related genes. The network displays the associations among 97 ion transport-related genes identified from GO term analysis of the PRPF-4 depletion vs. control dataset. Nodes represent individual genes, and edges represent five types of interaction evidence, as indicated by different colors. The network highlights two distinct, highly interconnected clusters comprising *twk* potassium channel genes and *del/deg* sodium channel genes.

**Supplementary Table 1.** List of alternative splicing events and genes showing significant changes following acute PRPF-4 depletion. Each row represents one alternative splicing event detected by rMATS. Columns: GeneID (WormBase gene identifier), FDR (false discovery rate), IncLevel_Control (exon inclusion level in control), IncLevel_IAA (exon inclusion level in IAA-treated), IncLevelDifference_IAA_Control (difference in exon inclusion level between IAA-treated and control), group (splicing types: SE, RI, MXE, A3SS, A5SS), status (direction of change in exon inclusion following IAA treatment: positive, increased inclusion; negative, decreased inclusion), WBGene (WormBase ID), Common_name (common gene name).

**Supplementary Table 2.** List of genes showing significantly increased expression following acute PRPF-4 depletion. Each row represents one gene. Data are from DESeq2 output. Columns: Gene (WormBase gene identifier), baseMean (mean of normalized counts across all samples), log2FoldChange (log2 fold change, IAA-treated vs control), lfcSE (standard error of log2 fold change), stat (Wald statistic), pvalue (raw p-value), padj (Benjamini-Hochberg-adjusted p-value), WBGene (WormBase ID), Common_name (common gene name), class (inc, increased expression following IAA treatment).

**Supplementary Table 3.** List of genes showing significantly decreased expression following acute PRPF-4 depletion. Each row represents one gene. Data are from DESeq2 output. Columns: Gene (WormBase gene identifier), baseMean (mean of normalized counts across all samples), log2FoldChange (log2 fold change, IAA-treated vs control), lfcSE (standard error of log2 fold change), stat (Wald statistic), pvalue (raw p-value), padj (Benjamini-Hochberg-adjusted p-value), WBGene (WormBase ID), Common_name (common gene name), class (dec, decreased expression following IAA treatment).

**Supplementary Table 4.** Alternative splicing events identified among genes associated with enriched GO terms. DESeq2 differential gene expression profiles and identified alternative splicing events are shown. Worksheet tab 1 lists cilia-related genes (n = 38), tab 2 lists ion transport-related genes (n = 97), and tab 3 lists splicing-related genes (n = 57). NA indicates that no alternative splicing events were identified for the corresponding gene. Splicing categories: SE, skipped exon; RI, retained intron; MXE, mutually exclusive exons; A5SS, alternative 5′ splice site; and A3SS, alternative 3′ splice site. Splicing status (positive, negative, not_sig, or NA) indicates the significance and direction of alternative splicing changes. Positive and negative indicate significantly increased or decreased alternative splicing, respectively, not_sig indicates no significant alternative splicing change, and NA indicates that no alternative splicing event was identified.

**Supplementary Table 5.** Alternative splicing events identified among differentially expressed genes. Significantly differentially expressed genes identified by DESeq2 and significantly differentially spliced events are shown. Splicing categories: SE, skipped exon; RI, retained intron; MXE, mutually exclusive exons; A5SS, alternative 5′ splice site; and A3SS, alternative 3′ splice site. Splicing status (positive or negative) indicates significantly increased or decreased alternative splicing, respectively, whereas RNA-seq status (inc or dec) indicates significantly increased or decreased gene expression, respectively.

## Notes

### Competing Interest Statement

The authors have declared no competing interest.

### Summary of Updates

Figure 3 and 4 revised. New Supplemental files added. Manuscript revised accordingly.

## References

Alahari, S. K., H. Schmidt and N. F. Kaufer, 1993 The fission yeast prp4+ gene involved in pre-mRNA splicing codes for a predicted serine/threonine kinase and is essential for growth. Nucleic Acids Res 21: 4079–4083.

Andrews, N. M., L. Gerten, A. Edwards, M. Brennan, Q. Reynolds et al., 2025 DiI Dye-Filling as a Simple and Inexpensive Tool to Visualize Ciliated Sensory Neurons in C. elegans. J Vis Exp.

Bacaj, T., Y. Lu and S. Shaham, 2008 The conserved proteins CHE-12 and DYF-11 are required for sensory cilium function in Caenorhabditis elegans. Genetics 178: 989–1002.

Baralle, F. E., and J. Giudice, 2017 Alternative splicing as a regulator of development and tissue identity. Nat Rev Mol Cell Biol 18: 437–451.

Cho, Y. S., J. Zhu, S. Li, B. Wang, Y. Han et al., 2018 Regulation of Yki/Yap subcellular localization and Hpo signaling by a nuclear kinase PRP4K. Nat Commun 9: 1657.

Dellaire, G., E. M. Makarov, J. J. Cowger, D. Longman, H. G. Sutherland et al., 2002 Mammalian PRP4 kinase copurifies and interacts with components of both the U5 snRNP and the N-CoR deacetylase complexes. Mol Cell Biol 22: 5141–5156.

Dobin, A., C. A. Davis, F. Schlesinger, J. Drenkow, C. Zaleski et al., 2013 STAR: ultrafast universal RNA-seq aligner. Bioinformatics 29: 15–21.

Eckert, D., N. Andree, A. Razanau, S. Zock-Emmenthal, M. Lutzelberger et al., 2016 Prp4 Kinase Grants the License to Splice: Control of Weak Splice Sites during Spliceosome Activation. PLoS Genet 12: e1005768.

Felix Krueger, F. J., Phil Ewels, Ebrahim Afyounian, Michael Weinstein, Benjamin Schuster-Boeckler, Gert Hulselmans, & sclamons, 2023 FelixKrueger/TrimGalore: v0.6.10 - add default decompression path, path (Version 0.6.10) [Computer software]. Zenodo. 10.5281/zenodo.7598955.

Fu, X. D., and M. Ares, Jr., 2014 Context-dependent control of alternative splicing by RNA-binding proteins. Nat Rev Genet 15: 689–701.

Gao, X., Q. Jin, C. Jiang, Y. Li, C. Li et al., 2016 FgPrp4 Kinase Is Important for Spliceosome B-Complex Activation and Splicing Efficiency in Fusarium graminearum. PLoS Genet 12: e1005973.

Habib, E. B., S. Mathavarajah and G. Dellaire, 2022 Tinker, Tailor, Tumour Suppressor: The Many Functions of PRP4K. Front Genet 13: 839963.

Hyung, D., J. Kim, S. Y. Cho and C. Park, 2018 ASpedia: a comprehensive encyclopedia of human alternative splicing. Nucleic Acids Res 46: D58–D63.

Kanno, T., P. Venhuizen, T. N. Wen, W. D. Lin, P. Chiou et al., 2018 PRP4KA, a Putative Spliceosomal Protein Kinase, Is Important for Alternative Splicing and Development in Arabidopsis thaliana. Genetics 210: 1267–1285.

Kew, C., W. Huang, J. Fischer, R. Ganesan, N. Robinson et al., 2020 Evolutionarily conserved regulation of immunity by the splicing factor RNP-6/PUF60. Elife 9.

Koterniak, B., P. P. Pilaka, X. Gracida, L. M. Schneider, I. Pritisanac et al., 2020 Global regulatory features of alternative splicing across tissues and within the nervous system of C. elegans. Genome Res 30: 1766–1780.

Liao, Y., G. K. Smyth and W. Shi, 2014 featureCounts: an efficient general purpose program for assigning sequence reads to genomic features. Bioinformatics 30: 923–930.

Lutzelberger, M., C. A. Bottner, W. Schwelnus, S. Zock-Emmenthal, A. Razanau et al., 2010 The N-terminus of Prp1 (Prp6/U5-102 K) is essential for spliceosome activation in vivo. Nucleic Acids Res 38: 1610–1622.

Matera, A. G., and Z. Wang, 2014 A day in the life of the spliceosome. Nat Rev Mol Cell Biol 15: 108–121.

Mathavarajah, S., S. Chipurupalli, E. B. Habib, W. D. Kim, M. M. Aoki et al., 2025 The evolutionarily conserved PRP4K-CHMP4B/vps32 splicing circuit regulates autophagy. Cell Rep 44: 115870.

Mazin, P. V., P. Khaitovich, M. Cardoso-Moreira and H. Kaessmann, 2021 Alternative splicing during mammalian organ development. Nat Genet 53: 925–934.

Moore, B. T., J. M. Jordan and L. R. Baugh, 2013 WormSizer: high-throughput analysis of nematode size and shape. PLoS One 8: e57142.

Neumuller, R. A., C. Richter, A. Fischer, M. Novatchkova, K. G. Neumuller et al., 2011 Genome-wide analysis of self-renewal in Drosophila neural stem cells by transgenic RNAi. Cell Stem Cell 8: 580–593.

Nilsen, T. W., and B. R. Graveley, 2010 Expansion of the eukaryotic proteome by alternative splicing. Nature 463: 457–463.

Nishimura, K., T. Fukagawa, H. Takisawa, T. Kakimoto and M. Kanemaki, 2009 An auxin-based degron system for the rapid depletion of proteins in nonplant cells. Nat Methods 6: 917–922.

Niu, L., K. Murugasen, S. Hanggodo, S. Ferdousy, L. Zhu et al., 2026 A scaffold attachment factor PHM-2 regulates synaptic transmission through SLO-2 potassium channel in C. elegans. PLoS Genet 22: e1011962.

Pan, Q., O. Shai, L. J. Lee, B. J. Frey and B. J. Blencowe, 2008 Deep surveying of alternative splicing complexity in the human transcriptome by high-throughput sequencing. Nat Genet 40: 1413–1415.

Power, K. M., J. S. Akella, A. Gu, J. D. Walsh, S. Bellotti et al., 2020 Mutation of NEKL-4/NEK10 and TTLL genes suppress neuronal ciliary degeneration caused by loss of CCPP-1 deglutamylase function. PLoS Genet 16: e1009052.

Power, K. M., K. C. Nguyen, A. Silva, S. Singh, D. H. Hall et al., 2024 NEKL-4 regulates microtubule stability and mitochondrial health in ciliated neurons. J Cell Biol 223.

Ragle, J. M., S. Katzman, T. F. Akers, S. Barberan-Soler and A. M. Zahler, 2015 Coordinated tissue-specific regulation of adjacent alternative 3’ splice sites in C. elegans. Genome Res 25: 982–994.

Robertson, S., and R. Lin, 2015 The Maternal-to-Zygotic Transition in C. elegans. Curr Top Dev Biol 113: 1–42.

Schneider, M., H. H. Hsiao, C. L. Will, R. Giet, H. Urlaub et al., 2010 Human PRP4 kinase is required for stable tri-snRNP association during spliceosomal B complex formation. Nat Struct Mol Biol 17: 216–221.

Schwelnus, W., K. Richert, F. Opitz, T. Gross, Y. Habara et al., 2001 Fission yeast Prp4p kinase regulates pre-mRNA splicing by phosphorylating a non-SR-splicing factor. EMBO Rep 2: 35–41.

Scotti, M. M., and M. S. Swanson, 2016 RNA mis-splicing in disease. Nat Rev Genet 17: 19–32.

Shen, S., J. W. Park, J. Huang, K. A. Dittmar, Z. X. Lu et al., 2012 MATS: a Bayesian framework for flexible detection of differential alternative splicing from RNA-Seq data. Nucleic Acids Res 40: e61.

Wang, S. J., and Z. W. Wang, 2013 Track-a-worm, an open-source system for quantitative assessment of C. elegans locomotory and bending behavior. PLoS One 8: e69653.

Wang, Z., and C. B. Burge, 2008 Splicing regulation: from a parts list of regulatory elements to an integrated splicing code. RNA 14: 802–813.

Warde-Farley, D., S. L. Donaldson, O. Comes, K. Zuberi, R. Badrawi et al., 2010 The GeneMANIA prediction server: biological network integration for gene prioritization and predicting gene function. Nucleic Acids Res 38: W214–220.

Watabe, E., S. Ono and H. Kuroyanagi, 2018 Alternative splicing of the Caenorhabditis elegans lev-11 tropomyosin gene is regulated in a tissue-specific manner. Cytoskeleton (Hoboken) 75: 427–436.

Weinreb, A., E. Varol, A. Barrett, R. M. McWhirter, S. R. Taylor et al., 2025 Alternative splicing across the C. elegans nervous system. Nat Commun 16: 4508.

Will, C. L., and R. Luhrmann, 2011 Spliceosome structure and function. Cold Spring Harb Perspect Biol 3.

Xia, P., L. Zhou, J. Guan, W. Ding and Y. Liu, 2022 Splicing factor PRP-19 regulates mitochondrial stress response. Life Metab 1: 81–93.

Yoshida, K., and S. Ogawa, 2014 Splicing factor mutations and cancer. Wiley Interdiscip Rev RNA 5: 445–459.

Zhang, L., J. D. Ward, Z. Cheng and A. F. Dernburg, 2015 The auxin-inducible degradation (AID) system enables versatile conditional protein depletion in C. elegans. Development 142: 4374–4384.

