## Supplementary figures and images for "The splicing kinase PRPF-4 is required for somatic development and germline function in *C. elegans*"

### Supplementary Figure 1

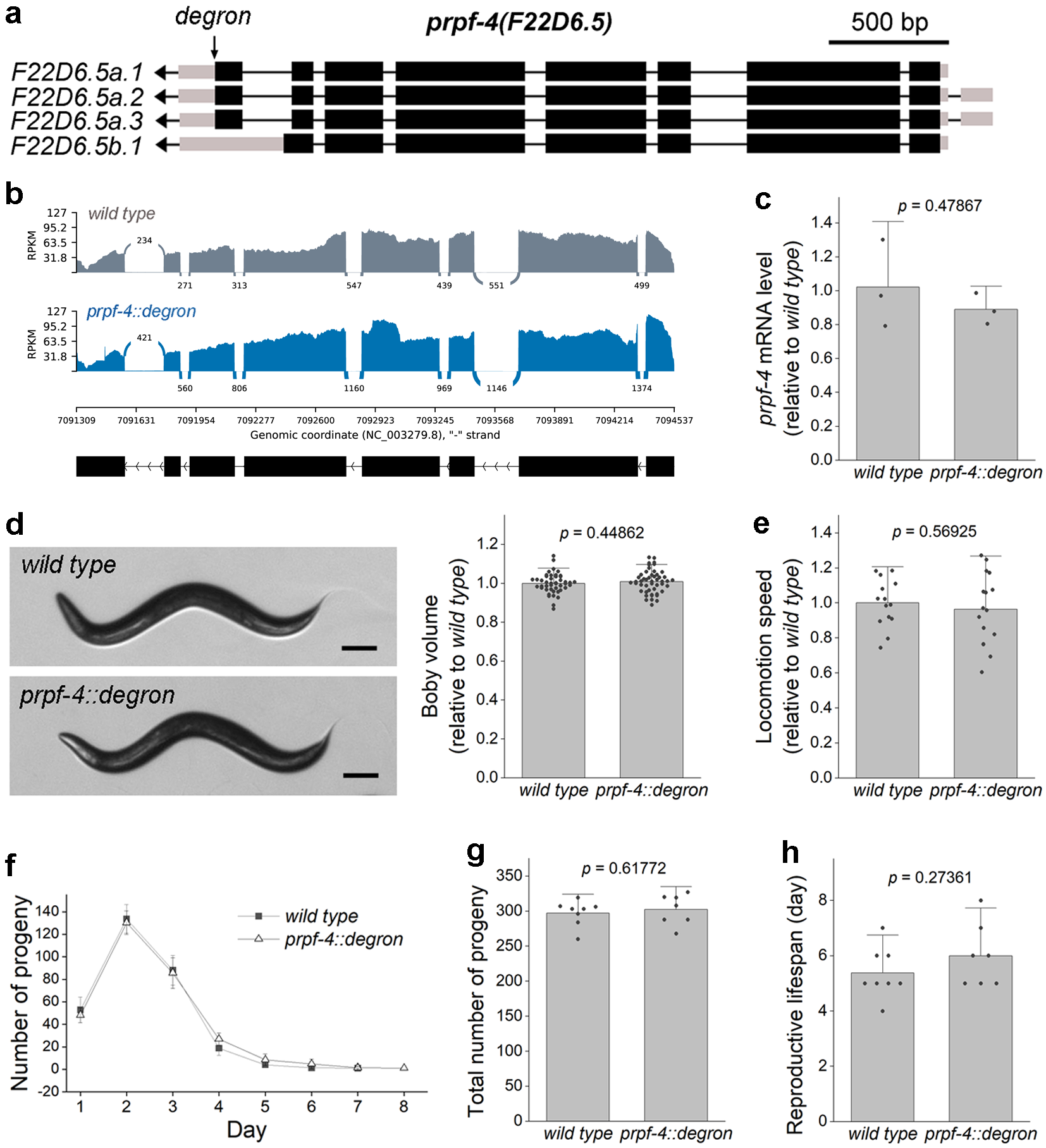

### Supplementary Figure 2

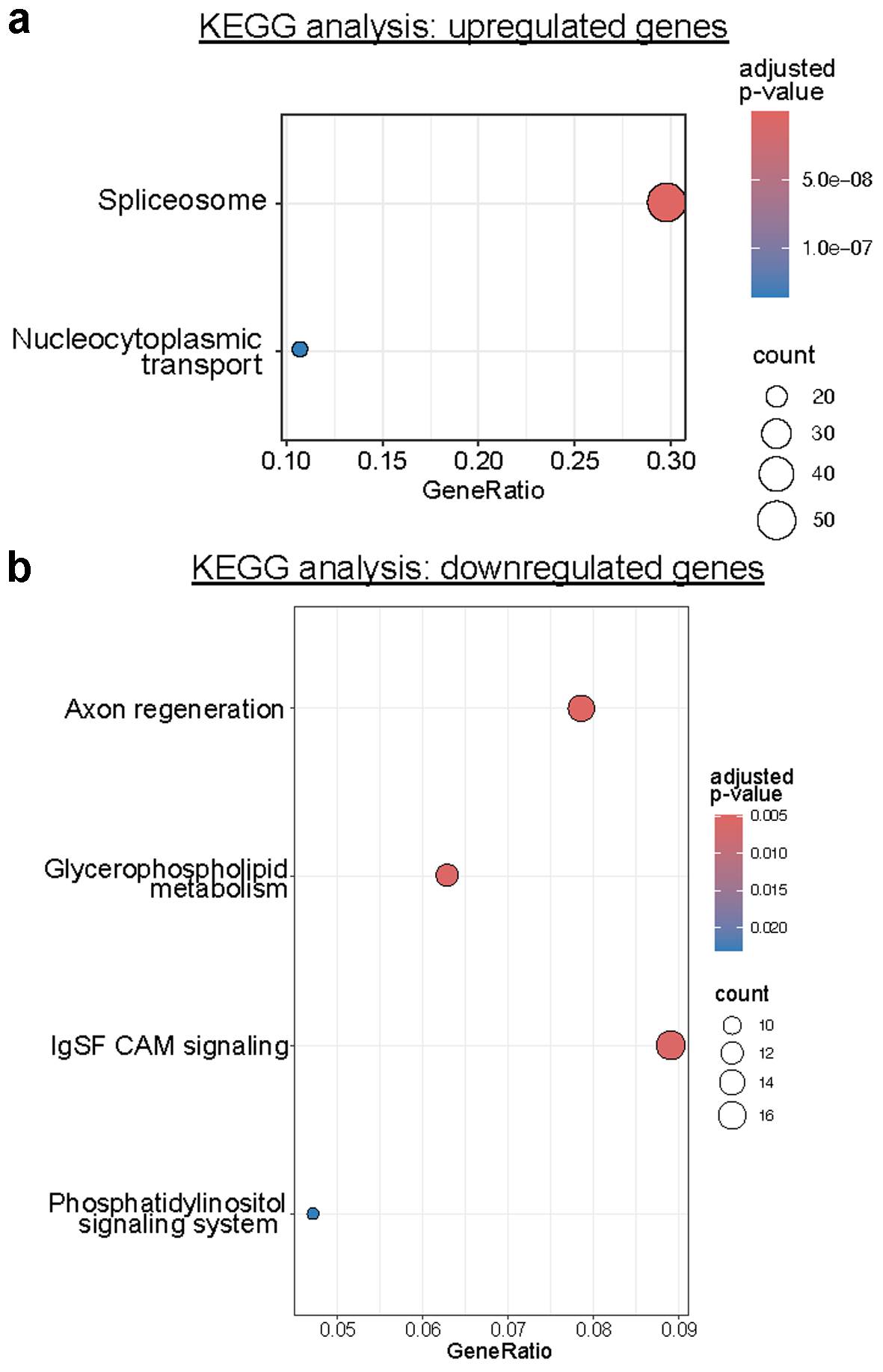

### Supplementary Figure 3

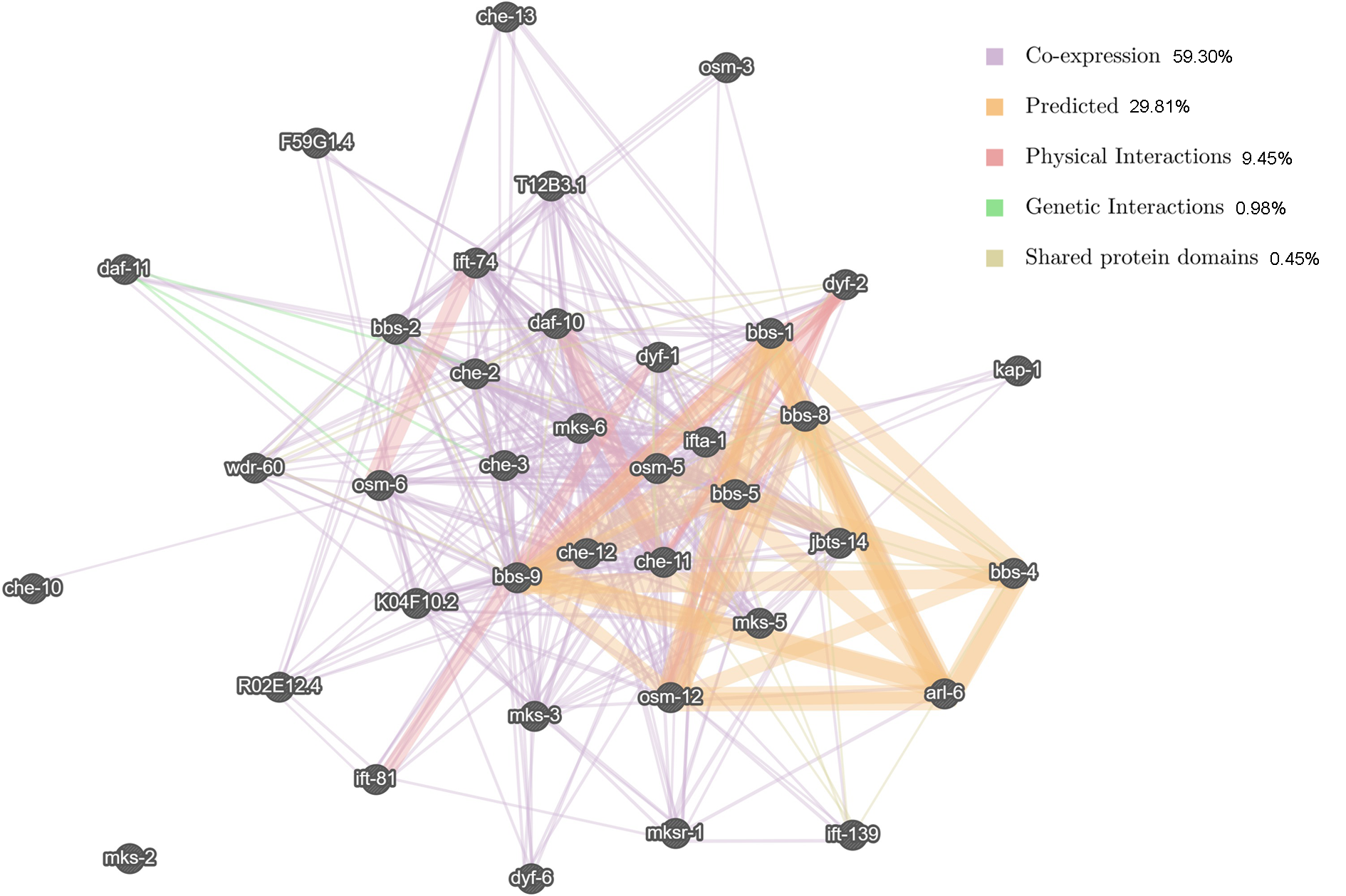

### Supplementary Figure 4

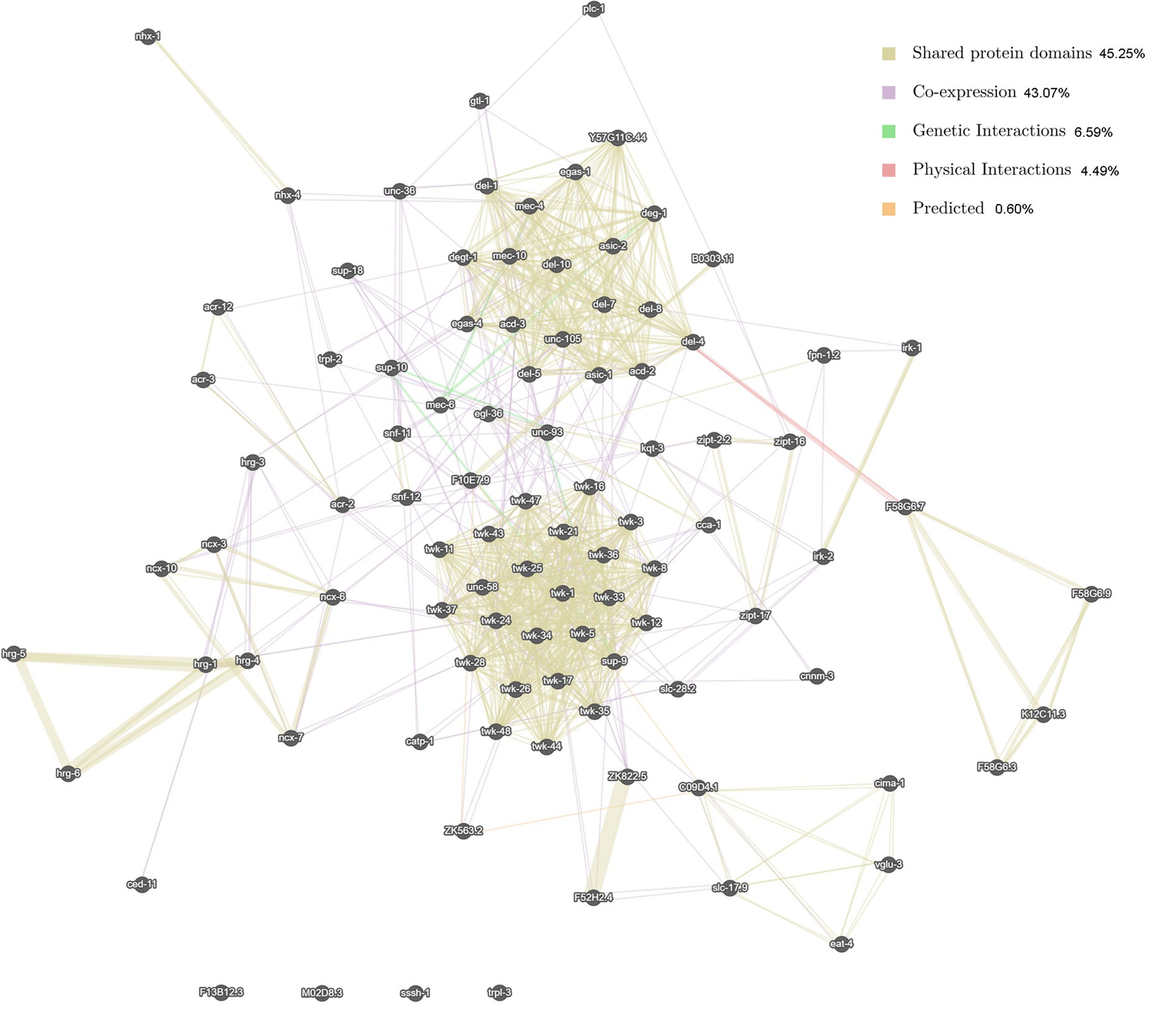
